# Complement biosynthesis in human brain: Insights from single-nucleus transcriptomics of hippocampus

**DOI:** 10.1101/2025.05.06.652374

**Authors:** Van Dien Nguyen, Matthew Bright, You Zhou, B. Paul Morgan, Wioleta M. Zelek

## Abstract

Complement is a key contributor to neuroinflammation, driving pathology in neurodegenerative diseases (NDDs); however, little is known about the source of complement in brain. Effective targeting of complement in NDDs requires understanding of its source; in particular, which brain cells express complement genes, how expression is regulated and how expression changes in disease.

To address this knowledge gap, we identified and integrated single-nucleus RNA sequencing (snRNA-seq) datasets from 398,097 nuclei across 48 hippocampal samples from non-demented brains to create a comprehensive transcriptomic atlas of complement gene expression ("complementome") in healthy brain. Expression levels of genes encoding complement components, receptors, and regulators were analysed across different brain cell types and the impact of sex and age on complement gene expression tested. To test impact of disease, we generated a further atlas from datasets comprising 11 non-demented and 12 Alzheimer’s disease (AD) patient hippocampi to assess changes in complement gene expression in AD.

All glial cells in the non-demented hippocampus expressed complement genes. *C1Q A/B/C* genes were exclusively expressed in microglia, while *C3* was highly expressed in microglia and astrocytes. Notably, *C3* expression defined a subset of pro-inflammatory microglia. Complement receptor encoding genes (*C3AR1*, *C5AR2*, *ITGAM*, *ITGAX*, *ITGB2*, *VSIG4*) were predominantly expressed in microglia, while neurons expressed a set of brain-specific putative receptors (*NPTX 1/2*, *NPTXR*, *NRP1*). Endothelial cells abundantly expressed key regulators (*CD46*, *CD55*, *CD59, CFH*), while neurons and OPCs expressed a set of putative regulators (*CSMD 1/2/3*, and *SUSD4*). Astrocytes abundantly expressed *CLU*. Cell type-specific gender differences included higher expression of *C1Q A/B/C*, *C1R*, *C1S* in female microglia and higher expression of *ITGAM* and *ITGAX* in male microglia. Generally higher complement gene expression was seen in older donors. Compared to controls, AD microglia showed higher expression of *C1QB* and *C3*, and AD astrocytes showed higher *C1S* and *C3* expression. Expression of *ITGAX* was elevated in AD microglia compared to controls, while many AD cell types demonstrated reduced expression of brain-specific putative receptors.

Defining the complementome in non-demented hippocampus provides a baseline for exploration of brain complement expression. Complement expression is upregulated in AD, likely contributing to neuroinflammation and highlighting its potential as a therapeutic target.

**Graphic abstract:** 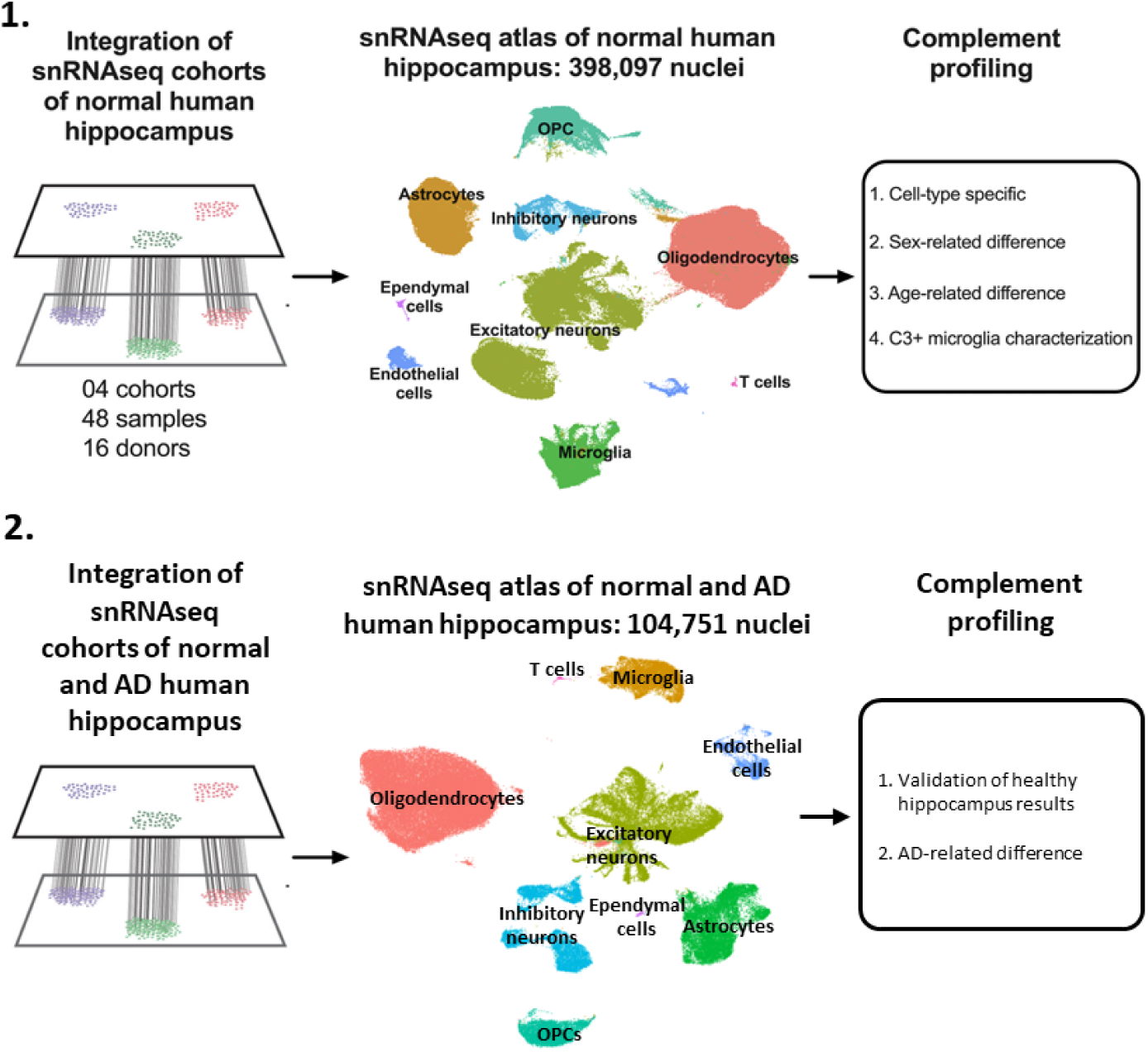

## Introduction

Complement, a central component of innate and adaptive immunity critical for immune defence and homeostasis, comprises a network of ∼60 proteins that collaborate to eliminate pathogens and toxic waste.^1^ Activation may be triggered via antibody-antigen complexes (classical pathway; CP) or directly on pathogen surfaces (lectin and alternative pathways; LP, AP); all activation pathways generate active fragments that either coat targets for phagocytic removal (opsonisation; C3b/iC3b, C4b) or activate cells to drive inflammation (anaphylaxis; C3a, C5a), activities mediated by complement receptors. The activation pathways converge on C3 cleavage and share a common terminal pathway (TP) that builds a pore, the membrane attack complex (MAC), in target cell membranes causing cell injury and inflammation. To restrict injury to self, host cells are protected by an armoury of regulators on membranes (CD46, CD55, CD59) and in the fluid phase (factor H, C4b binding protein, clusterin).

Despite its established role in immune defence, the relevance of complement in the central nervous system (CNS) has received sparse attention. In the brain, as in the periphery, complement plays protective roles; however, complement also influences brain development. During synaptic pruning, C1 binds synapses destined for elimination and activates the CP to deposit fragments of C4 and C3, marking these synapses for phagocytic elimination by microglia and astrocytes.^2–4^ Physiological pruning is crucial for healthy development and is implicated in forgetting in the healthy adult brain.^5^ Notably, the TP and MAC formation also contributes to synapse elimination.^6^ Several genes expressed primarily in brain encode putative complement regulators, including *SUSD4*, *SRPX2* and *CSMD1*; knock-out of these genes in mice is associated with abnormal social behaviours and aberrant neuronal morphology, further implicating complement in brain development.^7–10^

Complement dysregulation in brain has emerged as a key pathological mechanism in many neurodegenerative (NDDs) diseases, including Alzheimer’s disease (AD).^11,12^ Complement involvement extends beyond classical NDDs to neuropsychiatric and neurodevelopmental disorders, including schizophrenia (Sz)^3^ and autism spectrum disorder (ASD).^13,14^

In AD, multi-source evidence implicates complement in disease pathophysiology. Two of the top five hits in genome wide association studies are complement genes (*CLU* and *CR1*).^15^ Complement proteins including C1q, C3, and MAC, are markedly upregulated in vulnerable brain regions, decorating amyloid plaques.^16,17^ Work in mice has highlighted the role of complement in synapse loss associated with neurodegeneration; knockout of C1q (blocking CP) or C3 (blocking all pathways) ameliorated disease in AD models,^18–20^ while inhibition of MAC assembly via knockout of C6 or therapeutic blockade of C7 reduced synapse loss and rescued cognition in dementia models.^6,21^ Despite this abundance of evidence implicating complement in brain health and disease, insight into the sources of complement in the brain remains limited, posing challenges in understanding these conditions and developing effective therapeutics. Current knowledge, derived largely from animal models and *in vitro* investigations, could be significantly enhanced by leveraging the recent explosion of brain sequencing data.

To address this knowledge gap, we performed a comprehensive analysis of complement expression in non-demented and AD human hippocampus, a region critical for memory and learning, and among the earliest and most profoundly impacted regions in neurodegeneration.^22^ Through consolidation of single nuclear RNA-sequencing (snRNAseq) data acquired from 48 samples obtained from cognitively intact donors across 4 cohorts, we characterised the expression of 63 complement genes in nine cell types in the hippocampus. Our analysis revealed a complex cell type-specific landscape, demonstrating significant age- and sex-associated expression differences. Notably, we found that *C3* expression marks a subset of inflammatory microglia, suggesting a potential mechanism for early complement dysregulation which could drive excessive inflammation in the brain. To explore changes in complement expression in AD, we consolidated data from 23 additional hippocampus samples (12 AD, 11 healthy control) to generate a comparative atlas, providing a preliminary picture of complement gene expression changes indicative of complement dysregulation in AD hippocampus.

## Materials and Methods

### Data curation

The Gene Expression Omnibus (GEO, https://www.ncbi.nlm.nih.gov/geo/) data repository was searched for snRNAseq datasets of interest. We conducted a systematic search of hippocampal single-cell RNA sequencing data, incorporating both neurologically healthy controls and Alzheimer’s disease (AD) samples. The search strategy utilized key terms including “hippocampus”, “hippocampal”, “single nuclei”, “single cell”, “RNA sequencing”, “RNAseq”, and for the AD-specific datasets, “Alzheimer’s disease”, “AD”, “LOAD”, “EOAD”, “neurodegenerative disease”, and “dementia”. We further restricted our search to human (Homo sapiens) samples.

For the neurologically healthy controls, we focused exclusively on samples generated using 10x Genomics single-cell RNA sequencing to ensure consistency in sequencing technology. Healthy control samples from the following datasets were selected: GSE160189, GSE163737, GSE175814, GSE186538.^23–26^ In total, 48 hippocampal samples from 16 neurologically healthy donors were included, a full breakdown of samples and donor metadata can be seen in supplementary table 1 (Table S1).

For the AD cohort, because of the limited availability of AD single-cell RNA sequencing datasets we adopted a more inclusive approach without restricting to a single sequencing platform, selecting hippocampal samples from 23 donors (12 AD and 11 control) from datasets GSE175814, GSE138852 and GSE212606.^25,27,28^

All 10x Genomics generated datasets were downloaded in FASTQ format ready for further processing. For data generated by other sequencing platform (GSE212606), the counts matrix provided by the authors was downloaded for further analysis. The metadata available for the donors and samples can be found in supplementary table 2 (Table S2).

### Raw data processing

Quality control of raw reads was performed using FastQC (v0.11.9) (http://www.bioinformatics.babraham.ac.uk/projects/fastqc). Samples were required to have a minimum Phred quality score of 30 for >80% of bases and <10% adapter content to pass quality control. TrimGalore (v0.6.6) was then employed to trim low-quality bases and remove adapter sequences, using a quality threshold of 20 and a minimum length of 20 bp for retained reads (https://zenodo.org/records/7598955). Trimmed reads were next aligned to the 10X human reference genome GRCh38 (ref-2020-A) using CellRanger count (v6.1.2)^29^ on a Hawk supercomputer (Cardiff Supercomputing Facility; https://portal.supercomputing.wales/). The CellRanger output was imported into R (v4.1.0) using Seurat (v4.0.3).^30^ A Seurat object was created for each sample, with genes detected in fewer than three cells excluded. Low-quality nuclei were removed based on the following criteria: nuclei with <2ND percentile or >98th percentile of detected genes range, or >5% mitochondrial reads. To identify and remove doublets from our snRNAseq data, the DoubletFinder algorithm implemented in R was employed with an expected proportion of homotypic doublets adjusted for individual samples.^31^ Cells identified as doublets were removed from downstream analyses. Data were then normalized using SCTransform with default parameters. The top 3000 variable features were identified using the ’vst’ method in Seurat. The effects of mitochondrial percentage and number of UMIs were removed using the SCTransform function.

### Data integration, dimension reduction and clustering

From this step, data for samples comprising the cognitively healthy hippocampus atlas and the healthy/AD comparison atlas were processed separately. For each, Harmony (v1.0) was used to integrate data across all samples, using donor identities and sequencing batch as integration variables with theta set to 2 and nclust to 50.^32^ For the healthy/AD atlas, sequencing platform was added as a variable for integration. Dimensionality reduction was performed using principal component analysis (PCA) on the top 3000 highly variable genes identified by SCTransform. The first 30 principal components were used for uniform manifold approximation and projection (UMAP) visualization (n.neighbors=30, min.dist=0.3). The integrated data were clustered using the Leiden algorithm with a resolution of 0.8.

### Cell annotation

Cluster-specific markers were identified using Seurat’s FindAllMarkers function with the Model-based Analysis of Single-cell Transcriptomics (MAST) algorithm. Genes with an adjusted *P* value < .05 and log2 fold-change > 0.25 were considered significant markers. SingleR (v1.6.1) with the Human Primary Cell Atlas was used as a reference for automated annotation, using ’fine’ labels for validation of the marker-based annotation.^33^ Cell types were additionally annotated based on the expression of known marker genes: *RBFOX3*, *SLC17A7* (excitatory neurons); *RBFOX3*, *GAD1*, *GAD2* (inhibitory neurons); *GFAP*, *AQP4*, *SLC1A2* (astrocytes); *CX3CR1*, *P2RY12*, *APBB1IP* (microglia); *MOG*, *MBP*, *MOBP*, *PLP1* (oligodendrocytes); *PDGFRA*, *CSPG4*, *OLIG1* (OPCs); *CLDN5*, *VWF*, *PECAM1*, *FLT1* (endothelial cells); *CFAP54*, *FOXJ1*, *RSPH1* (ependymal cells); and *TRBC2*, *CD3D*, *CD3E* (T cells). Both approaches were largely consistent with each other, where conflicting results were obtained, the results from manual annotation using known marker genes were used.

### Complement gene expression

We analysed the expression of 63 complement genes (Supplementary Table 1) across all identified cell types, categorized into pathways (CP, AP, LP, TP), and as components, regulators or receptors. Gene expression was quantified using normalized and log-transformed counts. Dotplots were generated using Seurat’s DotPlot function and heatmaps were created using GraphPad Prism 8.4. Differential expression analysis was performed using Seurat’s FindMarker function with the Wilcoxon Rank Sum test and min.pct = 0.001 on the data slot of the SCT assay. The ident.1 argument defined the test condition (e.g. male, AD and >80years) and ident.2 defining the control condition (e.g. female, control donors and 20-40years). Genes with a Bonferroni adjusted *P* value < .05 were considered significant.

### C3-expressing microglia characterisation

Microglia were classified as *C3*-expressing if they had a detectable *C3* read count. Of the 42,490 microglia analysed, 18,684 (44%) were classified as *C3*-expressing. A pseudobulk differential expression analysis using DESeq2 (v1.32.0)^34^ was then applied to identify genes with differential expression between *C3*-expressing and *C3*-non-expressing cells. Pseudobulk samples were created by summing raw counts for each condition within each donor. To remove potential cross-dataset batch effects, we labelled each dataset with a distinct batch identifier then incorporated and adjusted the downstream analyses for identifier effects. Specifically, the design was fitted as “∼Batch + condition”, in which Batch included the list of identifiers while condition consisted of different comparison groups. A Benjamini-Hochberg adjusted *P* value < 0.05 was considered significant. Subsequently, the gene list from the differential expression analysis results was ranked based on stat value and subjected to pathway analysis using fast gene set enrichment analysis (fgsea v1.28.0) with minSize=15, maxSize=500, and nperm=10000 in the setting and biological pathway references from MSigDB database (https://www.gsea-msigdb.org/gsea/msigdb).^35^ Pathways with a Benjamini-Hochberg adjusted *P* value < .05 were considered significantly enriched.

### Other statistical analyses and plotting

Statistical analyses and figure generation were performed by using R v4.1.0 (http://www.r-project.org/), and GraphPad Prism 8.4 for Mac OS (GraphPad Software, La Jolla, CA).

## RESULTS

### Integrated single nuclei atlas of normal human hippocampus

To generate the transcriptional atlas of the complement system in non-demented human hippocampus we collected, reprocessed, integrated, and analysed four publicly available snRNAseq datasets (GSE160189, GSE163737, GSE175814, GSE186538 from GEO database), profiling brain cells through a common computational pipeline (see Methods). The 48 samples included were from 16 neurologically healthy donors and derived from hippocampus, among the first brain regions to manifest pathology in NDDs.^36^ The median age of donors was 50.5 years (interquartile range [IQR], 44–71.5 years), and five of sixteen were female. A full breakdown of donor and sample metadata (age, sex, dataset) is provided in supplementary table 1. All selected studies used 10x Single Cell RNAseq technology for sequencing. Following pre-processing, a total of 398,097 high quality nuclei were retained for downstream analysis. We then identified nine distinct cell types (excitatory neurons, inhibitory neurons, astrocytes, microglia, oligodendrocytes, oligodendrocyte precursor cells (OPCs), endothelial cells, ependymal cells, T cells) within the dataset, classified in an unbiased manner using a combination of established canonical marker genes and the SingleR tool.

### Cell-type-specific expression of complement components

#### a) Activation pathways

We first used the annotated atlas to characterise the expression landscape of the complement activation pathways.

In the CP, genes encoding the components of C1q, *C1QA*, *C1QB*, and *C1QC*, were uniquely expressed in microglia (Fig. 1A,E). Endothelial cells were the main expressors of *C1R* and *C1S*, encoding the serine proteases C1r and C1s that bind C1q to form the C1 complex. *C1R* and *C1S* expression was also detected in a subset of astrocytes and ependymal cells. *C2,* encoding C2, the protease component of the CP C3 convertase (C4b2a), was predominantly expressed by neurons and microglia. Surprisingly, the genes encoding the C4A and C4B isotypes, *C4A* and *C4B* demonstrated different expression patterns, the former primarily expressed by astrocytes and the latter by ependymal cells. Compared to C1-encoding genes, expression levels of *C2*, *C4A* and *C4B* were low.

**Figure 1.**
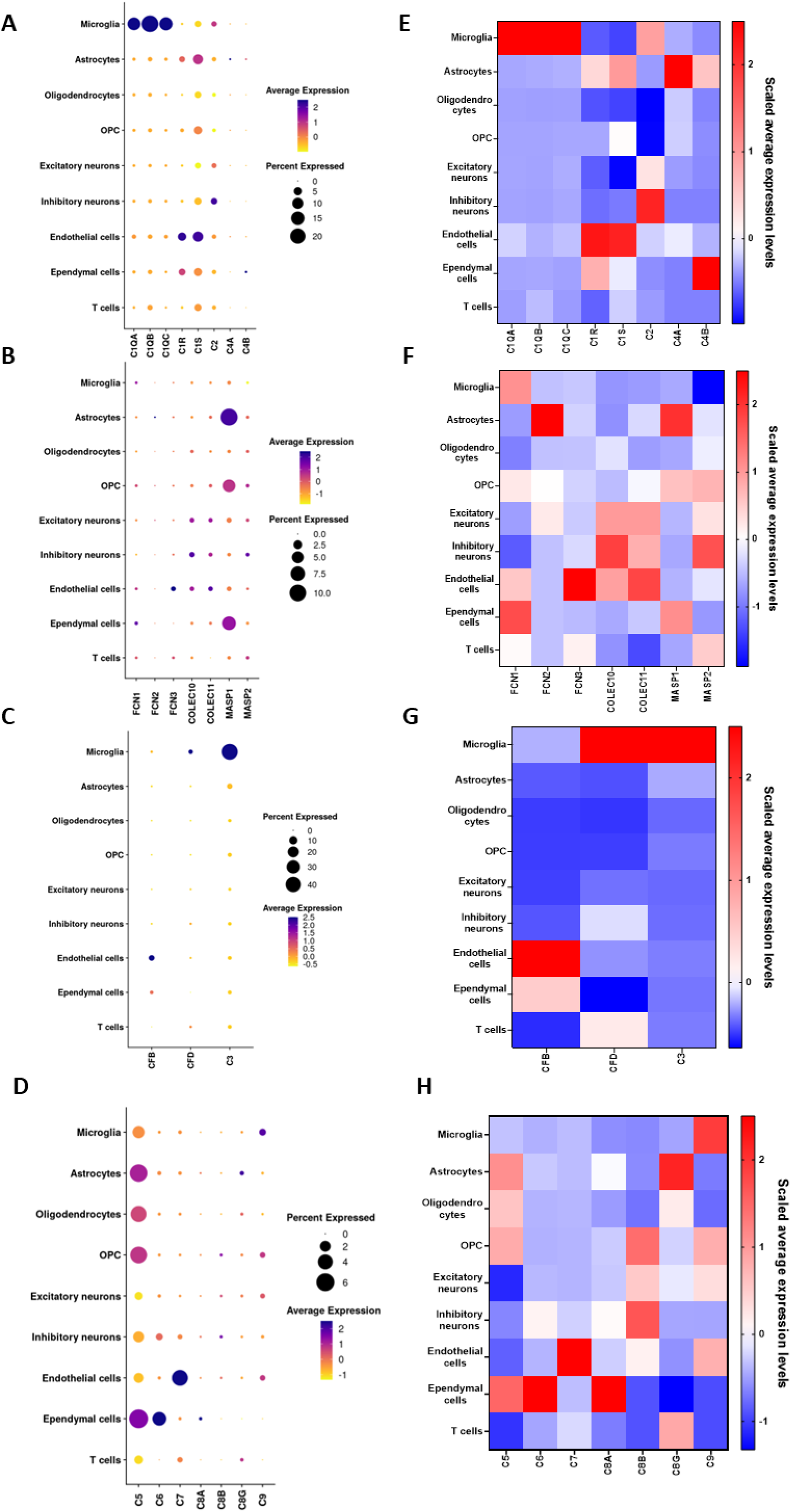
Expression profile of complement gene expression in hippocampus. **A-D.** Dot plots showing expression of complement CP, LP, AP, and TP genes in different cell types in the healthy human hippocampus. The size of the dots represents the percentage of cells expressing the gene, and colour intensity (from low in yellow to high in purple) represents the average expression level. In CP, exclusive expression of *C1QA*, *C1QB*, and *C1QC* in microglia was observed. In LP, *MASP1* was predominantly expressed in astrocytes, OPC and ependymal cells. In AP, expression of *CFB* was dominant in endothelial cells and *C3* was expressed predominantly in microglia. In TP, *C5* was widely expressed. **E-H.** Heatmaps showing expression levels of CP, LP, AP, and TP components across various brain cell types, including microglia, astrocytes, oligodendrocytes, oligodendrocyte precursor cells (OPC), excitatory neurons, inhibitory neurons, endothelial cells, and epithelial cells. Scaled expression levels are shown from low (blue) to high (red). In CP, microglia exhibited highly expressed *C1QA*, *C1QB*, and *C1QC*, absent in other cell types; endothelial cells showed notable expression of *C1S* and *C1R*, while inhibitory neurons demonstrated high expression of *C2*. In LP, endothelial cells expressed *MASP2*, *FCN3*, *COLEC10* and *COLEC11*, both inhibitory and excitatory neurons expressed predominantly *MASP2*, *COLEC10* and *COLEC11*, while ependymal cells showed moderate expression of *MASP1*. In AP endothelial cells exhibited significant expression of *CFB*. In TP endothelial cells expressed *C7*, while ependymal cells abundantly expressed *C5* and *C6*.

In the LP, genes encoding the pattern recognition molecules of the lectin pathway, *FCN1*, *FCN2*, *FCN3*, *COLEC10*, *COLEC11* and *MBL2* showed minimal or no detectable expression in the healthy hippocampus. Of the genes encoding the LP serine proteases, *MASP1* was expressed primarily by astrocytes with low expression in a small proportion of ependymal cells and OPCs while *MASP2* expression was not detected (Fig. 1B,F).

In the AP, *C3* (encoding C3, the core component of the complement system), was predominantly expressed in microglia (43.7% expressing *C3* gene) with low expression in astrocytes (3.9% expression) (Fig. 1C,G). *CFB* (encoding the AP serine protease factor B) was predominantly expressed in microglia while *CFD* (encoding factor D) was primarily expressed in endothelial cells (Fig. 1C,G).

#### b) Terminal pathway (TP)

Of the TP component-encoding genes, *C5* was the most widely expressed, detected consistently in all cell types, with astrocytes and ependymal cells the highest expressors followed by OPCs and oligodendrocytes (Fig. 1D,H). Expression of *C6* was highest in ependymal cells while *C7* expression was highest in endothelial cells. Expression of genes encoding other terminal complement components was low across all cell types, highest for *C9* in microglia, OPCs and excitatory neurons. Notably, *C8G*, encoding the gamma chain of C8, was abundantly expressed in a very small subset of astrocytes.

#### c) Complement receptors

The genes encoding receptors for C1q (*CALR, CD93, C1QBP*) were broadly expressed, albeit at low level, highest in endothelial cells, ependymal cells and neurons (Fig. 2A,B). Genes encoding the integrin receptors for C3 fragments CR3 and CR4 (*ITGAM, ITGAX* and *ITGB2)* were abundantly and exclusively (apart from low expression in T cells) expressed by microglia (Fig. 2A,B). The *CR1* gene was expressed abundantly in a small fraction of oligodendrocytes and microglia in healthy hippocampus, while *CR2* expression was restricted to T cells. The *VSIG4* gene, encoding the C3 fragment receptor CRIg (Complement Receptor of the Immunoglobulin Superfamily) was strongly and exclusively expressed in microglia (Fig. 2A,B). Genes encoding the anaphylatoxin receptors (*C3AR1, C5AR1, C5AR2*) were all expressed predominantly on microglia, although for *C5AR1* ependymal cells also showed strong expression. The genes encoding the neuropilins and neuronal pentraxins (*NRP1, NRP2, NPTX1, NPTX2, NPTXR*), putative complement receptors primarily expressed in brain, were widely expressed in brain cells, most abundantly in excitatory neurons (Fig.2A,B).

**Figure 2.**
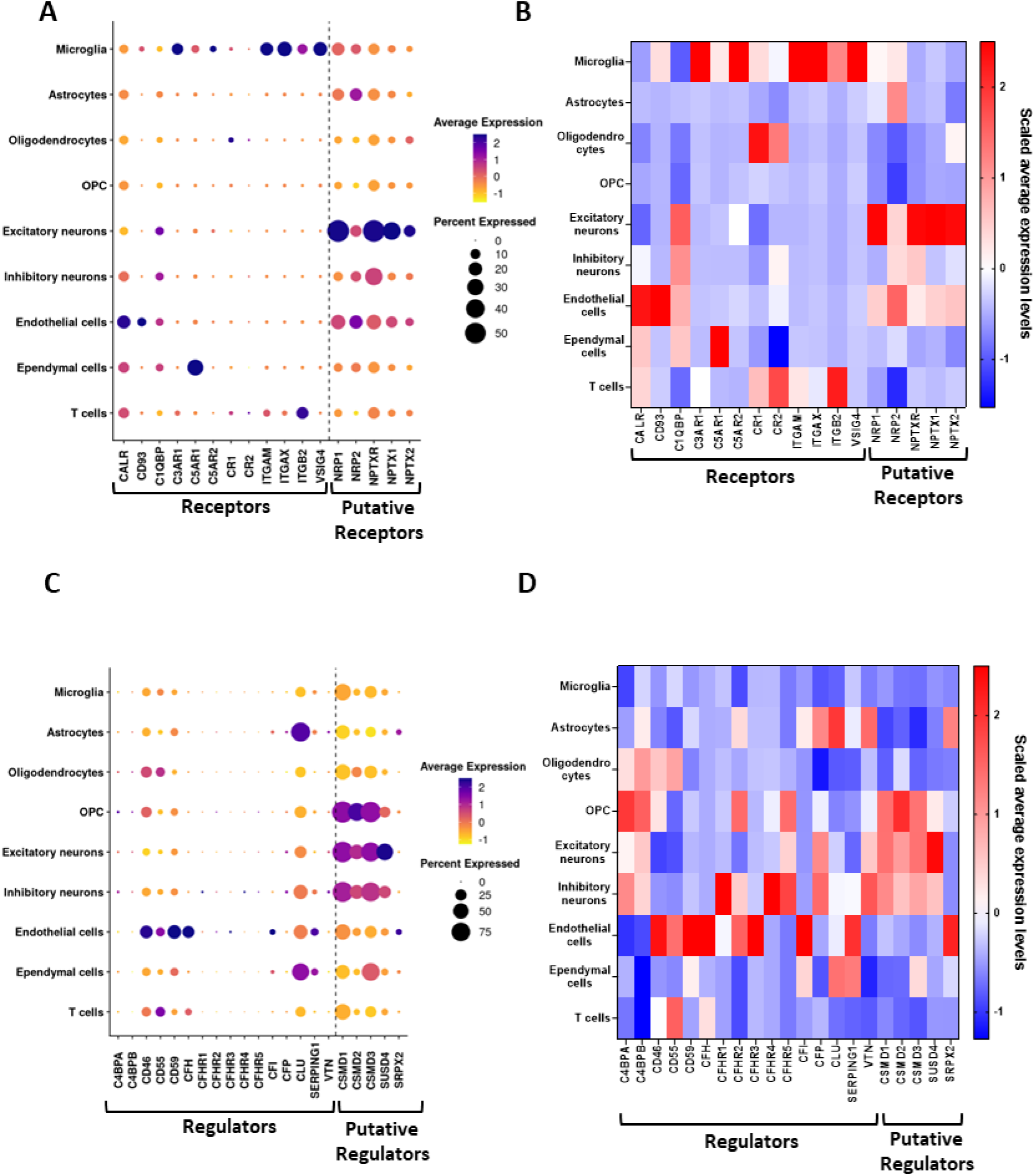
Expression profile of genes encoding complement regulators and receptors. **A.** Dot plot showing the expression of complement receptor genes across different cell types in the brain. The size of the dots represents the percentage of cells expressing the gene, and colour intensity (from low in yellow to high in purple) represents the average expression level. Excitatory neurons predominantly expressed *NPTX1*, *NPTX2*, and *NPTXR*; the other complement receptor genes, *ITGB2*, *ITGAX* and *VISG4*, were highly expressed in microglia, demonstrating cell-type-specific gene expression patterns. **B.** Heatmap displaying the expression levels of complement receptors. Scaled expression levels are represented from low (blue) to high (red). Microglia exhibited high expression of genes encoding complement receptors *C3AR1*, *C5AR1*, *ITGAM*, *ITGAX*, *ITGB2* and *VSIG4*; excitatory neurons showed expression of *NPTX1*, *NPTX2*, *NPTXR*, and *C1QBP*; endothelial cells displayed notable expression of *CALR* and *NRP1*. Oligodendrocytes expressed the highest levels of *CR1*, while ependymal cells expressed *C5AR1*. **C.** Dot plot depicting expression of genes involved in complement regulation in different cell types of the human healthy brain. All cell types expressed complement regulators, with *CSMD1, CSMD2, CSMD3* and *CLU* being predominantly expressed by most. Endothelial cells were the primary expressors of the complement membrane regulators *CD46, CD55, CD59* and *FH*. **D.** Heatmap showing expression levels of complement regulators. Scaled expression levels are displayed from low (blue) to high (red). Endothelial cells showed high expression of *CD46, CD55, CD59, CFH, CFI, SERPING1* and *SRPX2*, while astrocytes exhibited notable expression of *CLU* and *SRPX2*. Excitatory neurons predominantly expressed *CFP, CSMD1, CSMD2, CSMD3* and *SUSD4*. The *CSMD1-3* genes were also highly expressed by OPC. Inhibitory neurons also expressed *CFP* and *CSMD1-3*.

#### d) Complement regulators

Genes encoding the “classical” complement regulators (*CD46*, *CD55*, *CD59*, *CFH, SERPING1*) were most highly expressed in endothelial cells, but also in many other cell types at lower levels (Fig. 2C,D). *CLU*, encoding the terminal pathway regulator clusterin, was broadly expressed in all cell types, highest in astrocytes and ependymal cells, *CFI* was significantly expressed only in endothelial cells, while genes encoding other “classical” regulators, *VTN*, *CFP*, *CFH1 – CFH5*, were minimally expressed in brain cells (Fig. 2C,D). Of the putative regulators, genes encoding the CUB and Sushi Multiple Domain (CSMD; *CSMD1 – CSMD*3) and Sushi Domain Containing 4 (SUSD4; *SUSD4*) proteins were expressed by all cell types, most abundantly by OPCs and neurons, while *SRPX2* was expressed in a small subset of astrocytes and endothelial cells (Fig. 2C,D).

### Age- and sex-related differences in complement expression

To identify the impact of age and sex on complement expression in the healthy brain, samples were stratified by age (20-40, 41-60, 61-80, and >80 years) and sex. Peak expression in many For many complement pathway genes, expression increased with age, peaking in the 7^th^-8^th^ decades. Differential expression analysis of between the youngest and oldest age group revealed several significant changes in complement expression in the elderly (Fig. 3A). Notably, the abundantly expressed complement genes *C1QA*, *C1QB*, *C1QC* were all expressed at significantly higher levels in individuals aged over 80 years compared to those in the range of 20-40 years old with log2 fold changes of 1.66, 1.80, and 2.64 respectively (Fig. S1). *C2* expression was also increased significantly (1.69-fold; Fig. S1). In contrast, *C5* expression was significantly lower in over 80s (-0.56-fold change; Fig S1). Of the genes encoding complement receptors and regulators, the most significant increases in expression with age were observed for the brain-specific complement receptors (*NRP1*, *NRP2*, *NPTX1*, *NPTX2*, *NPTXR*) and regulators (*CSMD1*, *CSMD2*, *CSMD3*, *SUSD4*) (Fig. 3B; Fig. S1).

**Figure 3.**
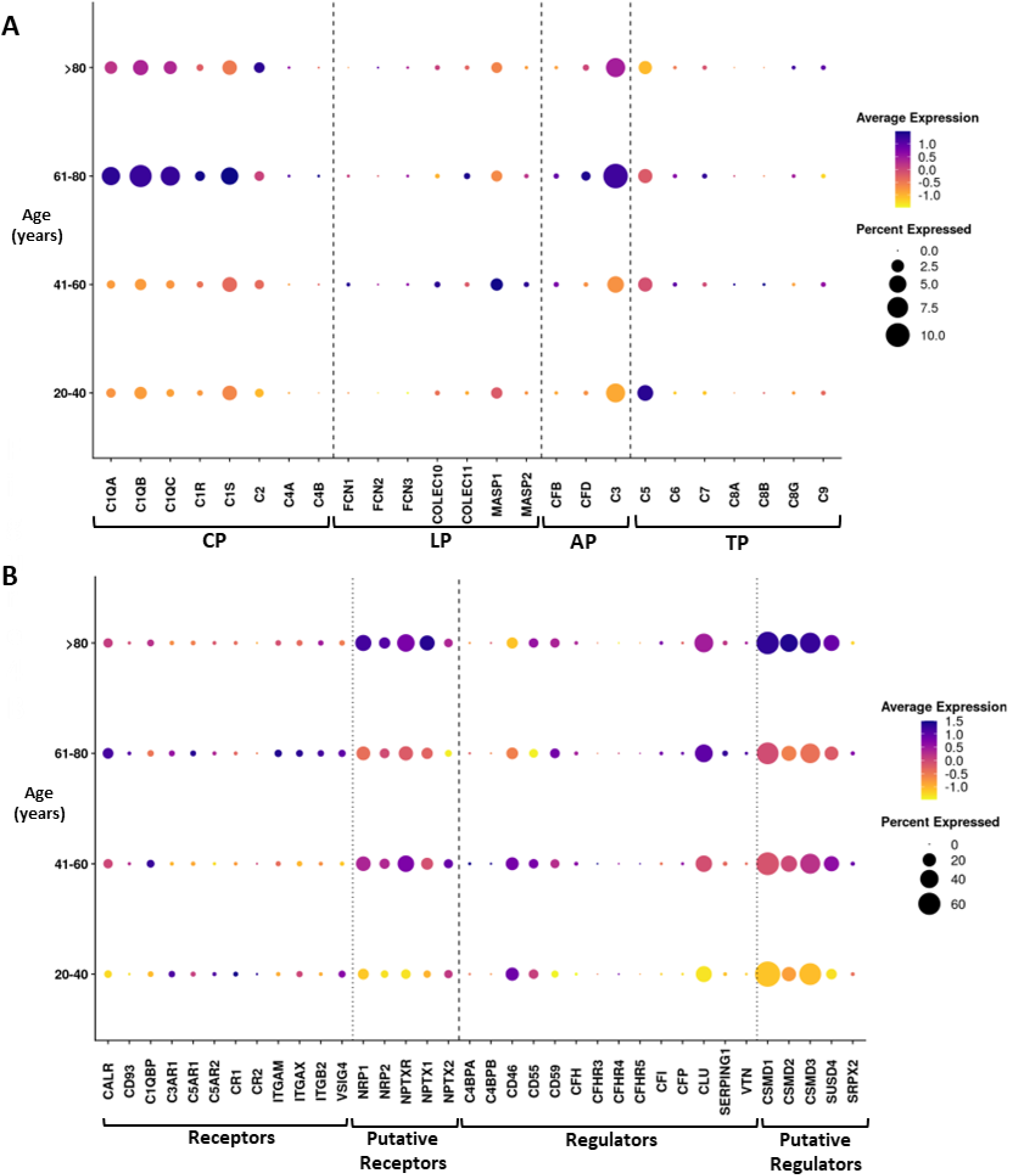
Age-related expression profiles of complement components, receptors, and regulators across brain cell types. The dot plots show the expression of various complement components, receptors, and regulatory proteins across brain cell types, stratified by age group. The size of the dots represents the percentage of cells expressing the gene, and colour intensity (from low in yellow to high in purple) represents the average expression level. **A.** Age-related changes in expression of complement component genes across CP, LP, AP, and TP; genes encoding components of the CP (*C1QA, C1QB, C1QC*, and *C1S*) and AP/TP (*C3, C5*) were more expressed in older age groups, peaking at 60 – 80 age group suggesting an increase in complement expression in aging brains. **B.** Age-related changes in genes encoding complement receptors and regulatory proteins, showing increased expression of *CLU, CSMD1-3, CFH, CFI, CD46* and *CD59* with age, peaking at >80 age group.

For most complement genes there was no difference in expression between the sexes. The exceptions were *C1QA*, *C1QB* and *C1QC* with expression in female microglia, astrocytes and oligodendrocytes all significantly greater than in male cells, more than two-fold in female astrocytes and microglia compared to male (Fig. S2A; Fig. S3A). *C1R* and *C1S* were also expressed at significantly higher levels in female astrocytes and endothelial cells compared to male (Fig. S2A; Fig. S3A). Female endothelial cells also had significantly higher expression of classical regulators *CD46*, *CD55* and *CD59,* compared to males, more than two-fold higher for *CD55* and *CD59* (Fig. S2B; Fig. S3B). The brain-specific receptors and regulators were expressed at higher levels in males across many cell types.

### Characterisation of C3-expressing microglia

Microglia were the predominant expressors of *C3* in healthy hippocampus with ∼44% of the total microglial population positive for *C3* (Fig. 1C; Fig. 4A). Comparative analysis demonstrated that the *C3*-positive microglia had significantly higher expression of other complement genes compared to *C3*-negative microglia; this included genes encoding classical pathway components (*C1QA/B/C*, *C1S*, *C2*), classical receptors (*CALR*, *C3AR1*, *C5AR1*, *ITGAM*, *ITGAX*, *ITGB2*, *VSIG4*), putative receptors (*NRP1/2*) and classical regulators (*CD46*, *CD55*, *CD59*) (Fig. 4B,C). Expression levels of some genes encoding regulators (*CLU*, *CSMD1/2/3*) and novel receptors (*NPTX1*, *NPTX2*, *NPTXR*) were lower in *C3*-expressing microglia (Fig. 4C). Pathway enrichment analysis confirmed the marked difference in complement profiles between *C3*-positive and *C3*-negative microglia with an upregulation of complement activation-related pathways (Fig. 4D). Pathway enrichment analysis also demonstrated that *C3*-positive microglia exhibited a higher inflammatory profile with significant enrichment in pathways related to myeloid cell activation, cytotoxicity, antigen processing/presentation and phagosome formation (Fig. 4E), innate and adaptive immunity (Fig. 4F), chemokine and cytokine production (Fig. 4G) and cell death (Fig. 4H).

**Figure 4.**
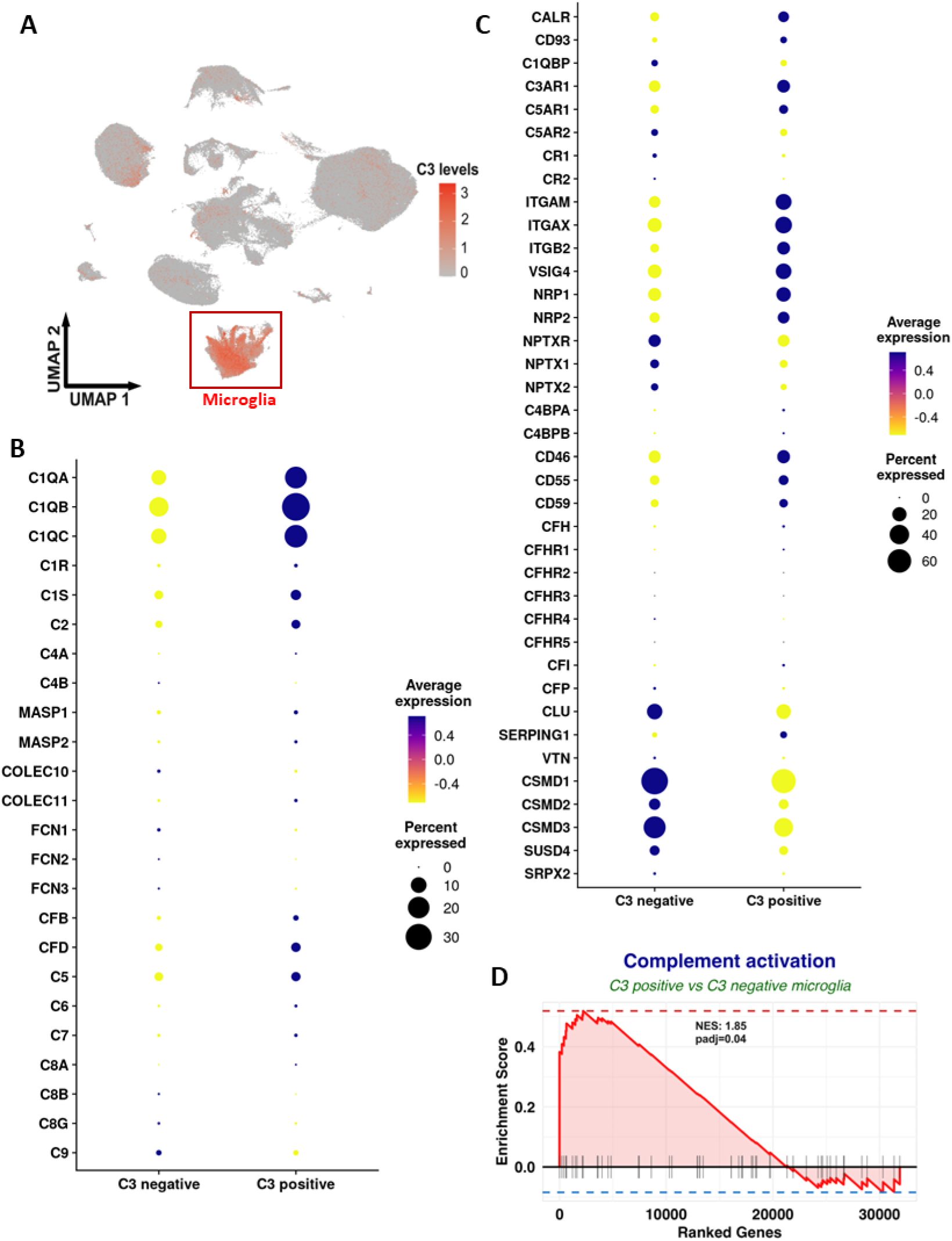
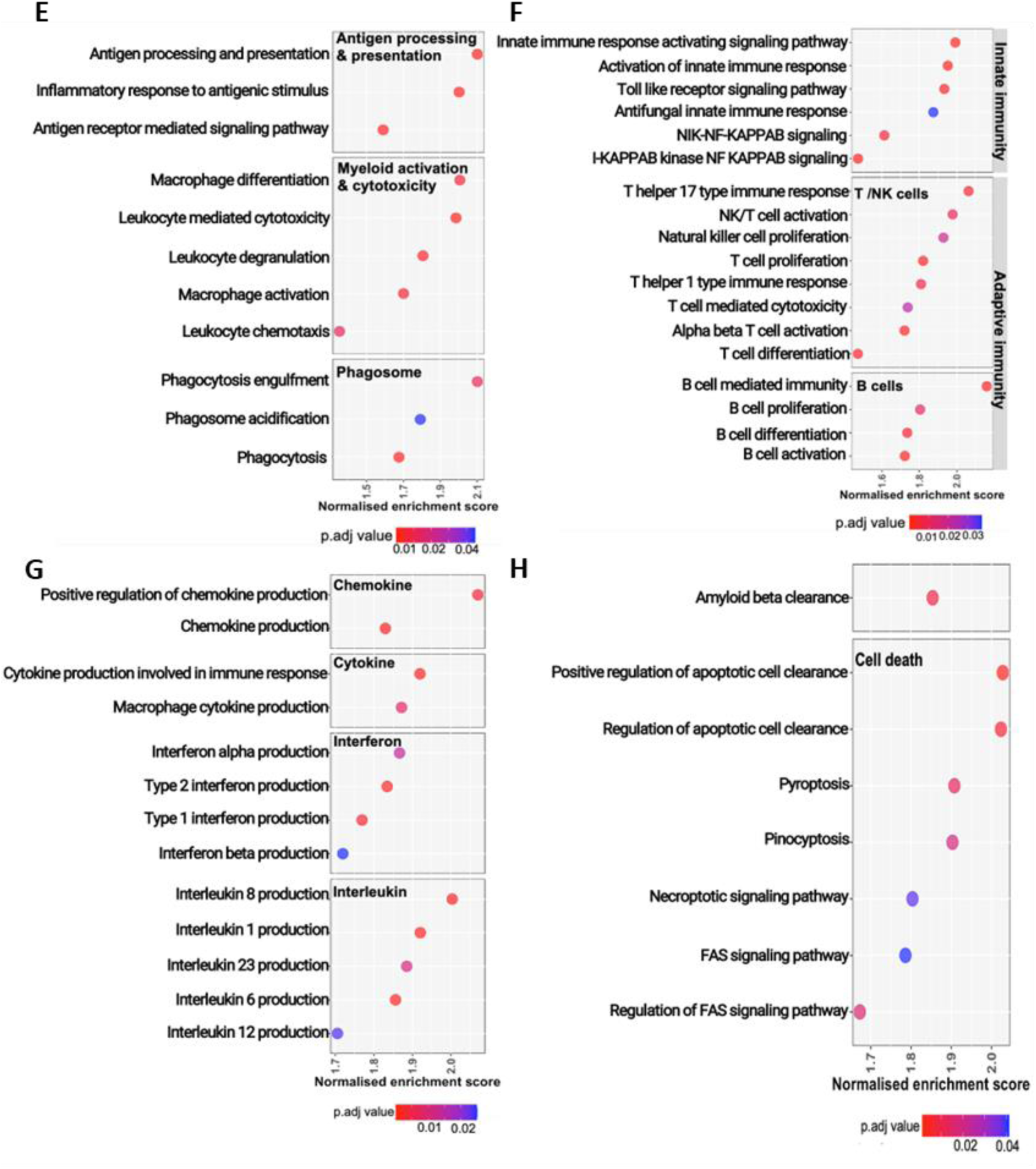
Expression profile of complement genes in *C3*-positive microglia. **A.** Uniform manifold approximation and projection (UMAP) plot illustrating *C3* expression in different cell types. The expression of *C3* is largely limited to microglia with a small population of *C3*-positive astrocytes. **B-C.** Dot plots showing the expression of various complement genes in *C3*-positive and *C3*-negative microglia in healthy hippocampus. Compared to *C3*-negative, *C3*-positive microglia showed elevated expression of genes encoding CP components *C1QA/B/C*, *C1S* and *C2*; receptors *CALR, C3AR1, C5AR1, ITGAM, ITGAX, ITGB2, VSIG4* and *NRP1-2*; regulators *CD46, CD55* and *CD59*. In contrast, *C3*-positive microglia exhibit reduced expression levels of complement regulators including *CLU, CSMD1-3*, and receptors *NPTX1, NPTX2, NPTXR*. **D.** Enrichment plot for complement activation in *C3*-positive versus *C3*-negative microglia; ranked list of genes with the enrichment score (y-axis) decreasing along the rank positions (x-axis) from left to right. Genes associated with complement activation are enriched at the beginning of the ranked list, indicating higher expression in *C3*-positive microglia. **E-H.** Dot plots of functional pathway analysis showing the normalised enrichment scores (x-axis) for various biological pathways related to immune and cellular functions in *C3*-positive microglia. Pathways are grouped by functional categories. Dot colours represent adjusted p-values, with more intense colours indicating greater statistical significance. Dot position reflects the enrichment score for the pathway. The most significant enrichment was observed in pathways related to immune activation, including “antigen processing & presentation” and “myeloid activation & cytotoxicity”, suggesting enhanced immune responses in *C3*-positive microglia. The “amyloid clearance” and “cell death” pathways also showed significant enrichment in association with apoptotic cell clearance and signalling, suggesting a potential link between immune activity and brain cell death.

### Integrated human atlas of the hippocampus in health and AD

Given the role of the complement system in AD progression and pathophysiology, we set out to identify changes in the expression of complement components in those with and without AD. To this end, we identified additional publicly available scRNA-seq datasets containing healthy and AD hippocampal samples (GSE138852, GSE175814, and GSE212606). Samples from neurologically healthy donors (n=11; nine were distinct from those used in the first analysis) and matched AD donors (n=12) were identified and processed through the pipeline used to generate the healthy atlas. A breakdown of the available metadata (age, sex, condition and dataset) can be found in supplementary table 2. The final atlas contained a total of 104,751 high quality nuclei (Fig. 5A). Nine distinct cell types (excitatory neurons, inhibitory neurons, astrocytes, microglia, oligodendrocytes, oligodendrocyte precursor cells (OPCs), endothelial cells, ependymal cells, T cells) were identified within the dataset using a combination of canonical marker gene expression and SingleR as for the first atlas.

**Figure 5.**
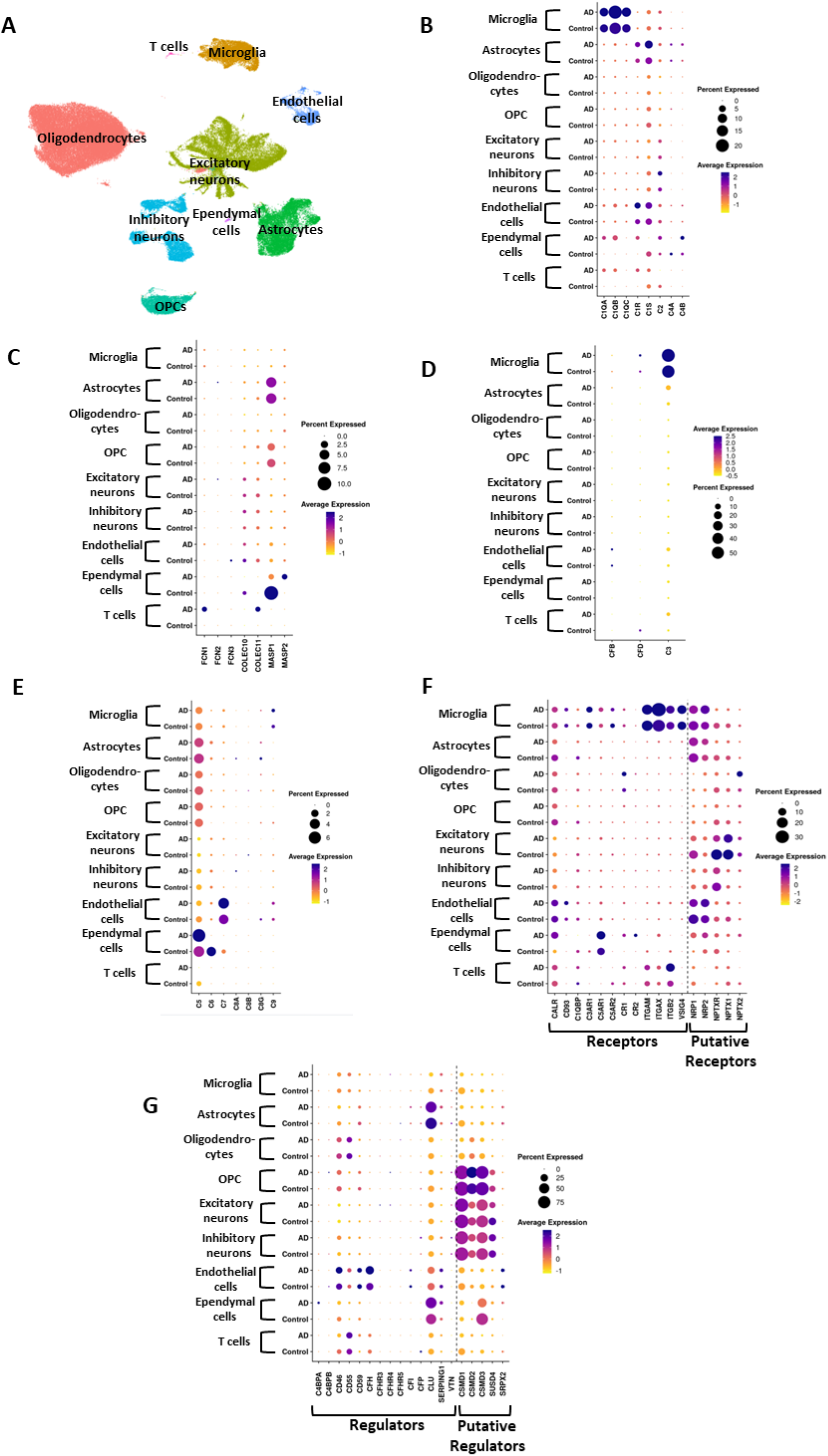
Complement gene expression profiles in non-AD and AD hippocampus. **A.** UMAP plot of brain single cell transcriptomes from 23 hippocampi (12 AD, 11 non-AD, 9 female, 14 male). From this set, expression of 63 complement genes in 9 cell types was measured and compared between non-AD and AD. **B-G.** Dot plots showing expression of complement CP, LP, AP, TP, receptor and regulator genes in different cell types; microglia, astrocytes, oligodendrocytes, oligodendrocyte precursor cells (OPC), excitatory neurons, inhibitory neurons, endothelial cells, ependymal cells and T cells of human hippocampi in health and AD. The size of the dots represents the percentage of cells expressing the gene, and colour intensity (from low in yellow to high in purple) represents the average expression level. In CP **(B)**, increased expression of *C1QA, C1QB*, and *C1QC* was observed in AD compared to non-AD microglia; *C1R* and *C1S* expression was increased in AD astrocytes and endothelial cells. In LP **(C)**, little change in expression was seen in AD. In AP **(D)**, *C3* expression was increased in AD in several cell types, predominantly microglia and astrocytes; FD was also upregulated in AD. In TP **(E)**, *C5* was upregulated in ependymal cells; *C7* expression was increased in endothelial cells. Among the complement receptors **(F)**, AD microglia demonstrate higher expression of genes encoding the anaphylatoxin receptors *C3AR1*, *C5AR2*, and C3b/iC3b receptors *ITGAM, ITGAX* and *ITGB2*. *C5AR1* expression was elevated in AD ependymal cells and *CR1* was elevated in AD oligodendrocytes. Of the complement regulators **(G)**, AD endothelial cells had higher expression of the classical regulators *CD46, CD55, CD59* and *CFH*; *SERPING1* was expressed at higher levels in AD across several cell types.

### AD-related changes in complement gene expression in the hippocampus

We first confirmed that complement gene expression in the non-demented control hippocampus samples in the comparative atlas recapitulated those in the original atlas then explored changes in expression in AD. Cell-type specific expression of complement genes was largely similar in AD; however, expression levels were different for several genes as detailed below.

#### a) Activation pathways

In the CP, AD microglia remained the dominant expressors of the C1q genes (*C1QA*, *C1QB*, *C1QC*). The proportion of microglia expressing C1q genes increased in AD and expression of *C1QB* was significantly upregulated, with a log2-fold change of 0.49 in AD microglia compared to healthy control (Fig. 5B; Fig. S4A). Expression of *C1QA* and *C1QB*, but not *C1QC*, was also detected in endothelial, ependymal and T cells in AD, though the increased expression did not reach significance. *C1R,* and *C1S* expression was increased in AD astrocytes and endothelial cells, significantly higher for the latter (48% increase; Fig. S4A). *C4A* and *C4B* expression was increased in several cell-types in AD; however, these changes were not significant (Fig. 5B).

Genes encoding LP recognition molecules (*FCN1*, *FCN2*, *FCN3*, *COLEC10*, *COLEC11*, *MBL2*) remained undetectable in AD hippocampus (Fig. 5C). Neither MASP1 nor MASP2 demonstrated significant changes in expression in AD though trends towards decreased expression were observed (Fig. S4).

For AP gene expression, *CFB* and *CFD* were not significantly different between AD and controls. As in healthy controls, microglia were the dominant expressors of *C3*, and expression was significantly increased in AD microglia; however, *C3* expression in AD astrocytes was much more significantly upregulated, increased 2.76-fold compared to controls (Fig. S4A). Endothelial, ependymal and T cells also showed increased *C3* expression in AD (Fig. 5D; Fig. S4A).

#### b) Terminal pathway (TP)

None of the genes encoding components of the TP showed significant changes in expression in AD compared to controls (Fig. 5E). Expression of *C5* and *C7* trended higher in AD but fell short of significance while other TP genes, expressed at low levels in the hippocampus, were unaltered.

#### c) Complement receptors

As in controls, the genes encoding the anaphylatoxin receptors (*C3AR1*, *C5AR1*, *C5AR2*) were primarily expressed in microglia in AD; significant upregulation of *C5AR1* was found in excitatory neurons and *C3AR1* trended towards increased expression in neurons and endothelial cells. *C5AR2* expression was decreased in many cell types. Expression of the genes encoding the integrin C3 fragment receptors *ITGAM* (encoding CR3, CD11b), *ITGAX* (encoding CR4, CD11c), *ITGB2* (encoding common CD18 chain) and *VSIG4* (encoding the Ig superfamily C3 fragment receptor CRIg) was essentially limited to microglia as in controls; *ITGAX* expression was significantly increased in AD microglia (54%; Fig. 5F; Fig. S4B). The gene encoding the C3b receptor CR1 (CD35; *CR1*) was expressed at low level on several cell types and significantly upregulated in oligodendrocytes in AD (log2 fold change of 0.60). Expression of the gene encoding the C3d receptor CR2 (CD21; *CR2*) was sparse and not significantly different between controls and AD (Fig. S4B). Expression of *NRP1*, encoding a putative C3/C4 fragment receptor, was significantly reduced in excitatory neurons and oligodendrocytes in AD (log2 fold changes of -1.42 and -2.65 respectively) (Fig. 5F). Genes encoding neuronal pentraxin receptors and neuronal pentraxin 1 (*NPTXR, NPTX1*) exhibited significantly lower expression in AD microglia, astrocytes, oligodendrocytes, OPCs and neurons.

#### d) Complement regulators

Expression patterns for most of the genes encoding “classical” complement regulators (*C4BP*, *FHRs3-5*, *FI*, *FP*) were similar between AD and non-AD hippocampus; the exceptions were *CD46* and *CD55*, significantly downregulated on oligodendrocytes and astrocytes in AD, and *CFH*, significantly upregulated in AD endothelial cells (Fig. 5G). Genes encoding the “putative” regulators, *SUSD4* and *CSMD1-3*, primarily expressed on neurons and OPCs, were significantly downregulated in AD (Fig. S4B).

## DISCUSSION

Evidence implicating complement in brain development and disease has accumulated over the last decade.^6,15–17,21,37,38^ Although these studies have identified critical roles of complement in brain health and disease, expression of complement proteins, regulators and receptors in brain has received scant attention. Understanding local production, activation and regulation of complement is essential for unravelling precisely how complement contributes to brain homeostasis and pathology. Early studies of complement expression in brain utilised cell lines and animal models, focussed on a handful of complement proteins and genes, and left large gaps in our understanding that need to be filled.^39–42^ Recent advances in single cell/nucleus sequencing technologies provide an opportunity to address this unmet need and gain an in-depth knowledge of local complement expression in brain.

We developed a comprehensive snRNA-seq expression atlas for complement genes in the non-demented human hippocampus, a region critical for learning and memory and impacted early in AD.^36^ The dataset comprised 398,098 nuclei, segregated into nine major cell types. We assessed 63 complement genes, encoding pathway components, regulators and receptors, revealing a diverse and intricate pattern of complement gene expression across brain cell types. Key findings include the demonstration that *C1QA/B/C*, encoding the three chains of the CP recognition moiety C1q, were abundantly and exclusively expressed in microglia, consistent with reports that microglial ablation eliminates C1q protein expression in rodent brain.^43,44^ The microglia-specific transcription factor PU.1, encoded by *SPI1* a known AD risk gene, modulates expression of *C1QA/C*, perhaps explaining the unique microglial expression of C1q.^45^ Others have implicated microglia-derived C1q in synaptic pruning during development^46,47^ and homeostasis in the aging brain.^48^ Microglial C1q directly activates other glia and provides a phagocytic signal.^4,49–51^ The genes encoding the C1q-associated proteases, *C1R/S*, were not expressed in microglia; endothelial cells (including pericytes in this analysis) showed highest expression with astrocytes and ependymal cells also significantly expressing both genes. The C1 complex components are thus made by different cell types, necessitating extracellular assembly, replicating the systemic situation where C1q is produced by macrophage/myeloid cells while C1r and C1s are synthesised in liver necessitating complex assembly in plasma.^52^ A recent preprint showed C1r/C1s expression in mouse brain choroid plexus (containing ependymal cells), further validating our findings.^53^ Expression of genes encoding other CP components (*C2*, *C4A, C4B*) was very low in non-demented hippocampus, suggesting limited baseline CP activity.

Expression of the LP pattern recognition molecules was extremely low or absent; absence of *MBL2* expression was supported by data from published brain atlases.^54,55^ The LP serine protease gene *MASP1* (alternatively spliced to yield MASP1, MASP3 and MAp44) was abundantly expressed in astrocytes and at lower levels in OPCs and ependymal cells. *MASP1* may have non-canonical roles in the developing brain; it was implicated in developmental neuronal migration in mice,^56^ and mutations in *MASP1* underpin the neurodevelopmental disorder 3-MC syndrome. ^57,58^

Expression of the AP-specific genes *CFB* and *CFD* was sparse, the former expressed at low levels by endothelial cells and the latter by microglia, suggesting low baseline AP activity in hippocampus. In contrast, expression of *C3,* encoding the core complement protein C3, was high in microglia with astrocytes and other cell types showing lower expression. The predominance of microglial expression of *C3* replicates Human Protein Atlas data (https://www.proteinatlas.org/).

Of the TP-specific genes, *C5* stood out, most abundant in ependymal cells but expressed by many cell types; expression of *C6* and *C7* was limited to ependymal cells and endothelial cells respectively while the genes encoding C8 and C9 were minimally expressed, except for a small subset of *C8G*-expressing astrocytes and a small *C9*-expressing microglial population.

Expression of *C5* may relate to non-canonical roles of C5a in brain homeostasis and response to injury, including roles in neural progenitor cell proliferation,^59^ brain homeostasis^60,61^ and synaptic plasticity.^62^ Low expression of other TP genes suggests that MAC formation is limited in the healthy brain.

Complement receptors were broadly but heterogeneously expressed across cell types. Genes encoding C1q receptors were highly expressed in endothelial cells and at low level in most brain cell types, providing a potential mechanism for the reported pleiotropic effects of C1q on neurons and glia.^48–51^ Anaphylatoxin receptor genes (*C3AR1, C5AR1, C5AR2*) were highly expressed in microglia, expected given their myeloid origins and likely critical contributing to complement-mediated microglial activation; however, by far the highest expression of *C5AR1* was in ependymal cells, a finding supported by data from Human Protein Atlas showing that choroid plexus has the highest brain expression of C5aR1. C5aR1 expression at this site may support the suggestion that choroid plexus is a “hotspot” for complement activation in brain.^63^ Genes encoding phagocytic C3 fragment receptors (CR3, CR4, CRIg; *ITGAM, ITGAX, ITGB2, VSIG4*) were all predominantly expressed by microglia, anticipated given that microglia are the brain professional phagocytes; astrocytes and T cells expressed these genes at low level with *ITGB2* a standout in T cells. Expression of *CR1,* encoding the C3b/C4b receptor, was highest in oligodendroglia and astrocytes, while *CR2* expression was minimal. Genes encoding neuropilins and neuronal pentraxins (*NRP1/2*, *NPTX1/2/R*), putative brain complement receptors, were abundantly expressed in neurons, highest in excitatory neurons, and at lower level in all other brain cell types. These proteins have been implicated in neuronal function and synaptic plasticity and shown to interact with complement proteins, although proof of complement receptor function is lacking.^64–67^

Genes encoding the membrane regulators (*CD46, CD55, CD59*) were expressed on all cell types, most abundantly on endothelial cells. Genes encoding the fluid-phase regulators were, in general, expressed at very low levels in hippocampus; the exception was *CLU,* broadly expressed and most abundant in astrocytes and ependymal cells, while endothelial cells expressed *CFH, CFI* and *SERPING1*. *CSMD1-3* and *SUSD4* were expressed in all cell types, highest in excitatory and inhibitory neurons and OPCs, while *SRPX2* was expressed in a small subset of astrocytes and endothelial cells. Of these, *CSMD1* has received the most attention; in mouse brain, *Csmd1* expression was found primarily in neurons and at synapses where it regulated complement activation.^10^

Microglia were the predominant *C3*-expressing cell type in healthy hippocampus; given the recent focus on astrocyte expression of *C3* as a marker of neurotoxic reactive (A1) astrocytes, upregulated in NDDs,^50^ we tested whether microglial *C3* expression also delineated functionally distinct populations. In non-demented hippocampus ∼44% of microglia expressed *C3;* these exhibited a pro-inflammatory phenotype with upregulation of complement components, receptors, and inflammatory mediators. These findings suggest that *C3* expression flags a functionally distinct microglial subset with enhanced pro-inflammatory and phagocytic capacity. Whether these changes reflect microglial priming reported in ageing brain and in NDDs remains unclear.^68,69^

Age stratification revealed an general increase in complement gene expression, particularly in individuals over 60 years old, aligning with the concept of brain "inflammaging"^70^ and implying increased complement dysregulation that might impact susceptibility to neurodegeneration in older individuals. Comparison of complement gene expression in females and males demonstrated a few significant differences, the most notable being higher *C1QA/B/C* expression in female microglia and higher *C1R/S* expression in female astrocytes and endothelia. Given the critical roles for C1q and C1 in developmental and pathological synaptic elimination, the increased expression of these genes might contribute to higher risk of AD in females.^71^

To explore changes in complement gene expression in AD, a separate snRNA-seq hippocampal dataset, comprising 104,751 nuclei from non-demented controls and matched AD samples, was collected, segregated into nine cell types and analysed through the established pipeline. Complement gene expression in controls closely replicated data from the original set. In comparison, the matched AD dataset showed differences in complement gene expression, including significantly upregulated expression of *C1QB*, *C1S*, and *C3*. Astrocyte expression of *C3* was significantly increased in AD, likely reflecting astrocyte activation to the *C3*-expressing A1 state.^50^ Expression of genes encoding ‘classical’ complement regulators was little changed in AD; however, genes encoding brain-specific putative complement regulators were widely downregulated in AD. Expression of genes encoding neuropilins and neuronal pentraxins was significantly downregulated in AD neurons. Neuropilins, key players in neurodevelopment,^64^ were implicated as complement receptors based on observed binding to C3d and C4d fragments, a tenuous association given the lack of any supporting functional evidence.^65^ Neuronal pentraxins bind C1q and influence microglial synaptic pruning^66,67^; reduced expression in AD may thus increase synapse loss. A better understanding of the function of these novel complement proteins will be essential for properly understanding the complement system in the brain.

Our snRNA-seq analyses provide a comprehensive reference map of complement gene expression in non-demented hippocampus, reveal a complex and tightly regulated landscape of complement gene expression, and give insight into changes in AD. Our focus was on hippocampus, but ongoing work will extend to other brain regions. Improved understanding of complement expression and dysregulation in the brain will help inform the development of targeted therapies for AD and other neurodegenerative diseases where complement is implicated.

## Data Availability

Information on the data underpinning this publication, including access details, can be found in the Cardiff University Research Data Repository at [DOI - TBC].

## Acknowledgements

We wish to honour the memory of Prof Philippe Gasque, whose pioneering work on complement expression in the brain and in brain cell lines during his time in Cardiff provided a crucial foundation for the development of the brain complement atlas. His dedication and contributions have left a lasting impact on the field, and his scientific legacy continues to inspire. We acknowledge the support of the Supercomputing Wales project, which is part-funded by the European Regional Development Fund (ERDF) via Welsh Government. We also acknowledge funding support from the UK Dementia Research Institute [UKDRI-3002] through UK DRI Ltd, principally funded by the Medical Research Council, The Moondance Foundation and Race Against Dementia.

## Supplementary Materials

**Supplementary Figure 1.**
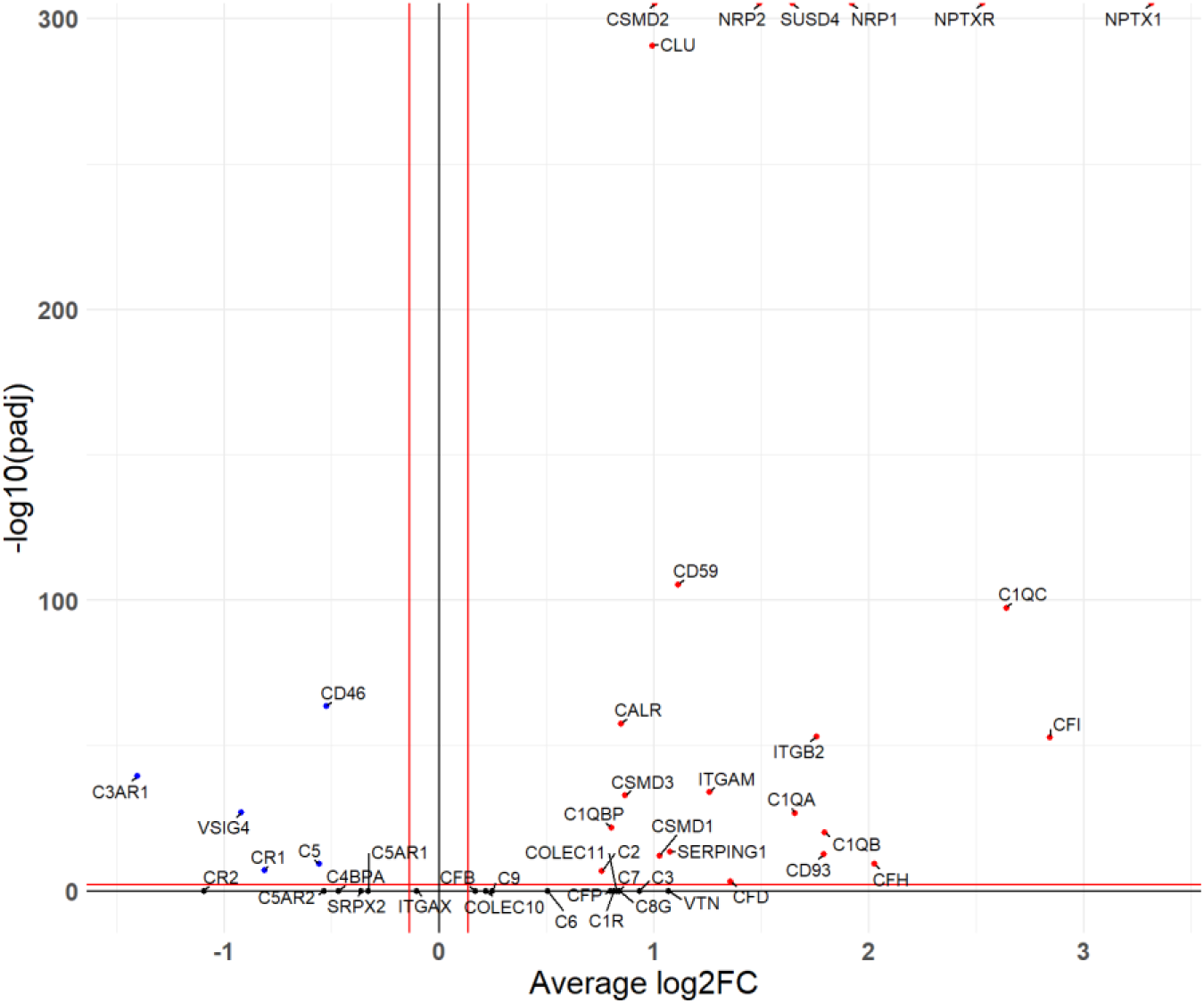
Differential expression analysis of complement genes in ageing. A volcano plot showing log2 fold change in expression on the x-axis and the -log10 of the Bonferroni adjusted *P*-value on the y-axis. Differential expression analysis was run comparing the 20-40-years age category to the >80-years category. *C1QA/B/C* and *C2* were all significantly upregulated in those >80 years old compared to 20-40 year olds. Many genes encoding putative complement regulators and receptors were also significantly upregulated in older donors.

**Supplementary Figure 2.**
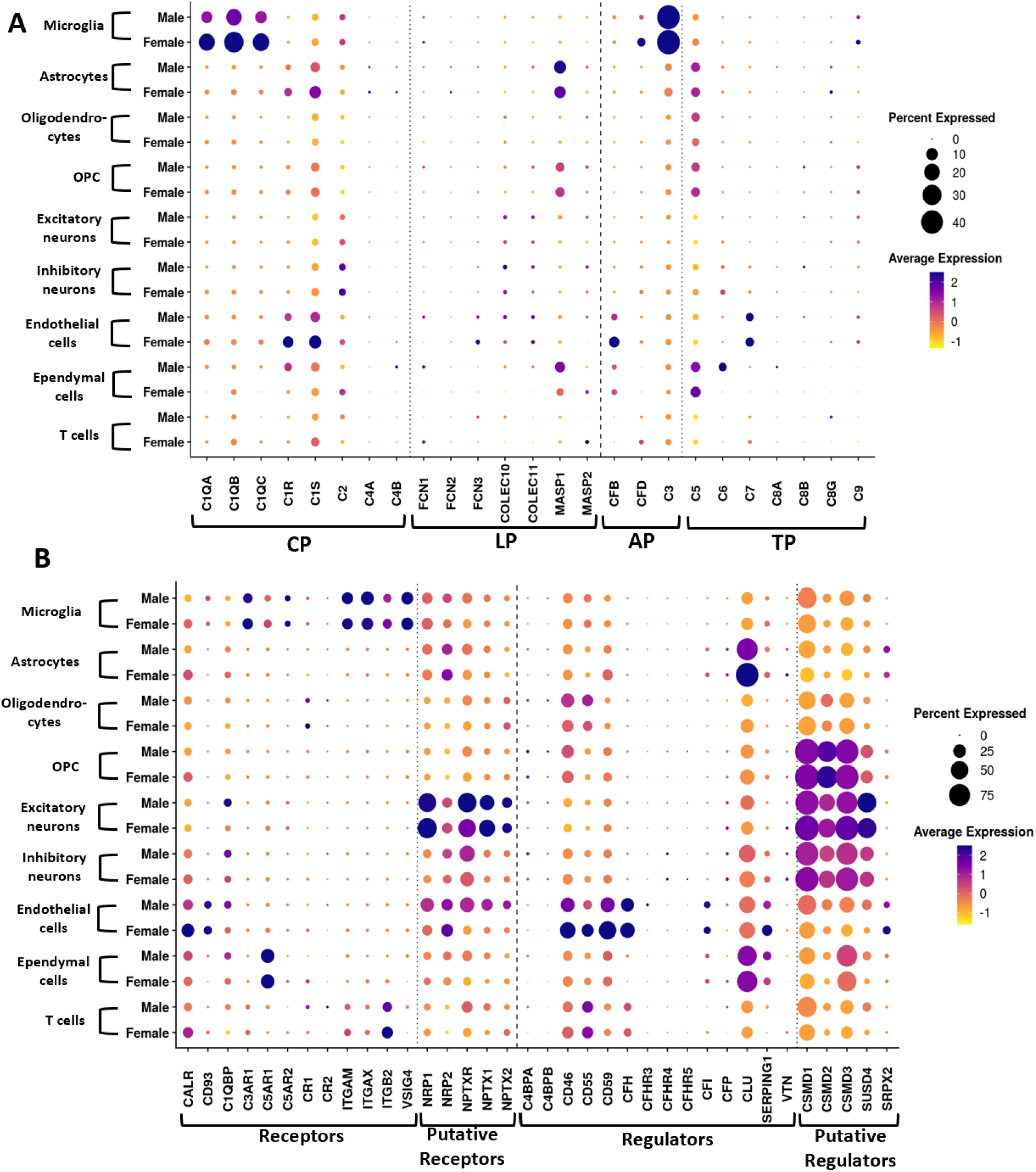
Sex-specific expression profiles of genes encoding complement components, receptors, and regulators across brain cell types. The dot plots illustrate expression in different brain cell types, separated by sex (donors: male, n = 12; female, n = 4), to highlight sex-based differences in complement expression. Each dot represents the percentage of cells expressing the gene, with larger dots indicating a higher proportion of expressing cells and colour intensity representing average expression levels (from low in yellow to high in purple). In (**A**), expression of genes encoding complement components across CP, LP, AP and TP is displayed for each cell type in male and female groups Notable differences include higher expression of *C1QA/B/C* in female microglia. In (**B**), expression of genes encoding complement receptors and regulatory proteins is shown divided by sex across brain cell types. Differences in expression patterns of several genes encoding complement regulators (*CFH, CD55, CD59, CLU*) were observed suggesting sex-based differences in complement regulation in brain cells.

**Supplementary Figure 3.**
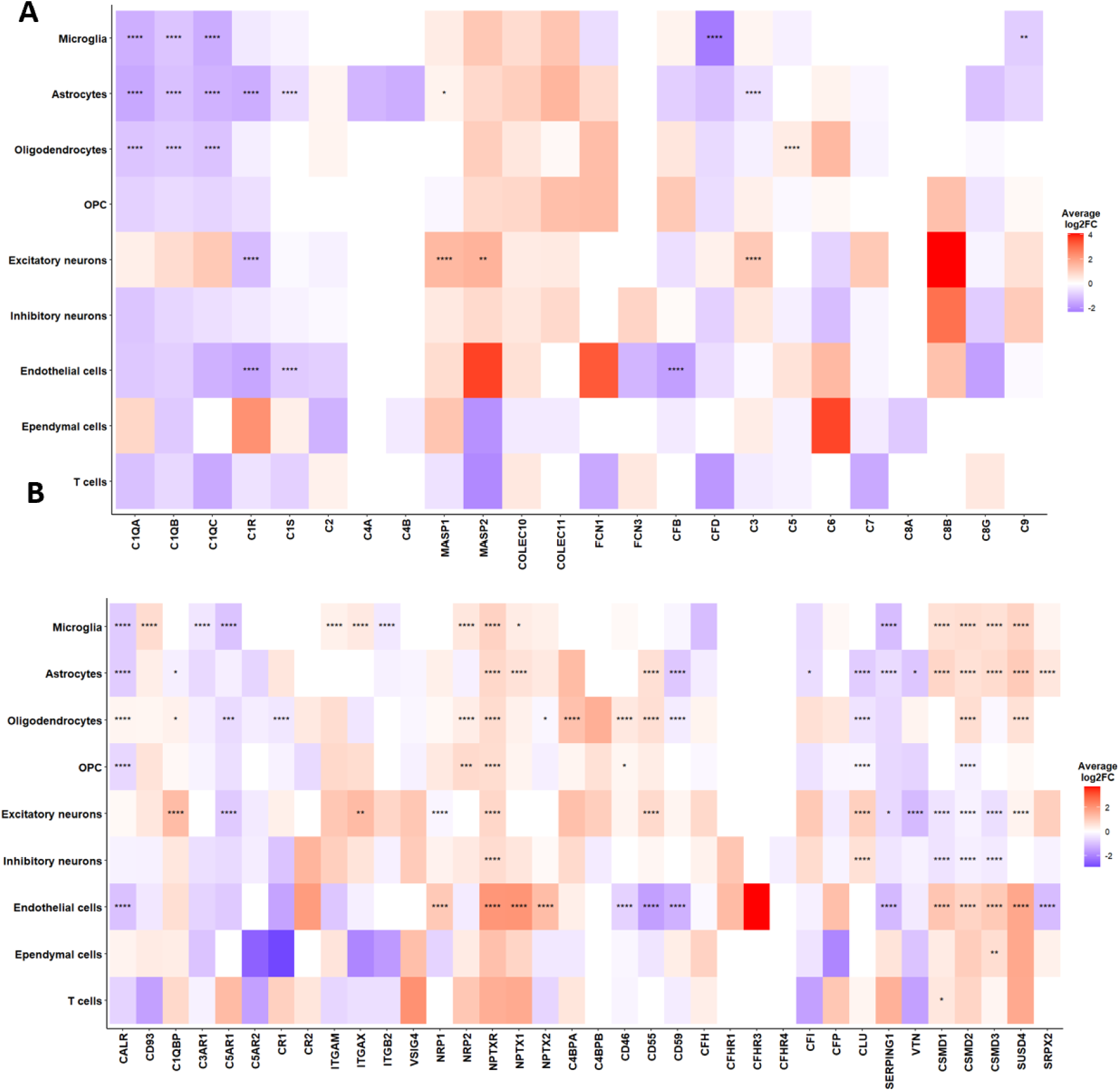
Differential expression of complement gene expression between males versus females within cell types. **A.** heatmap summarising the differences in complement expression between the sexes, within cell types. Differential expression analysis (DEA) was run in every cell type, the average log2 fold change is represented from low (blue) to high (red). A negative log2 fold change represents greater expression in female cells and a positive log2 fold change representing greater expression in male cells. Boxes are labelled based on their Bonferroni adjusted *P*-values (* = *P*<0.05, ** = *P*<0.01, *** = *P*<0.001, **** = *P*<0.0001). A. DEA summary for the genes of the complement pathways in the different cell types identified in the hippocampus. Females exhibited significantly greater expression of *C1QA/B/C* and *C1R* and *C1S*. Females also expressed *CFB* and *CFD* at greater levels than males. **B.** DEA summary for the complement receptor and regulator genes in the different cell types identified in the hippocampus. Male microglia expressed significantly greater levels of *ITGAM* and *ITGAX*. Males expressed significantly greater levels of several putative complement receptors including *NPTXR* and *NPTX1*. Of the putative regulators, female neurons expressed the CSMD genes at greater levels. Female endothelial cells express *CD46*, *CD55* and *CD59* at greater levels than male endothelial cells.

**Supplementary Figure 4.**
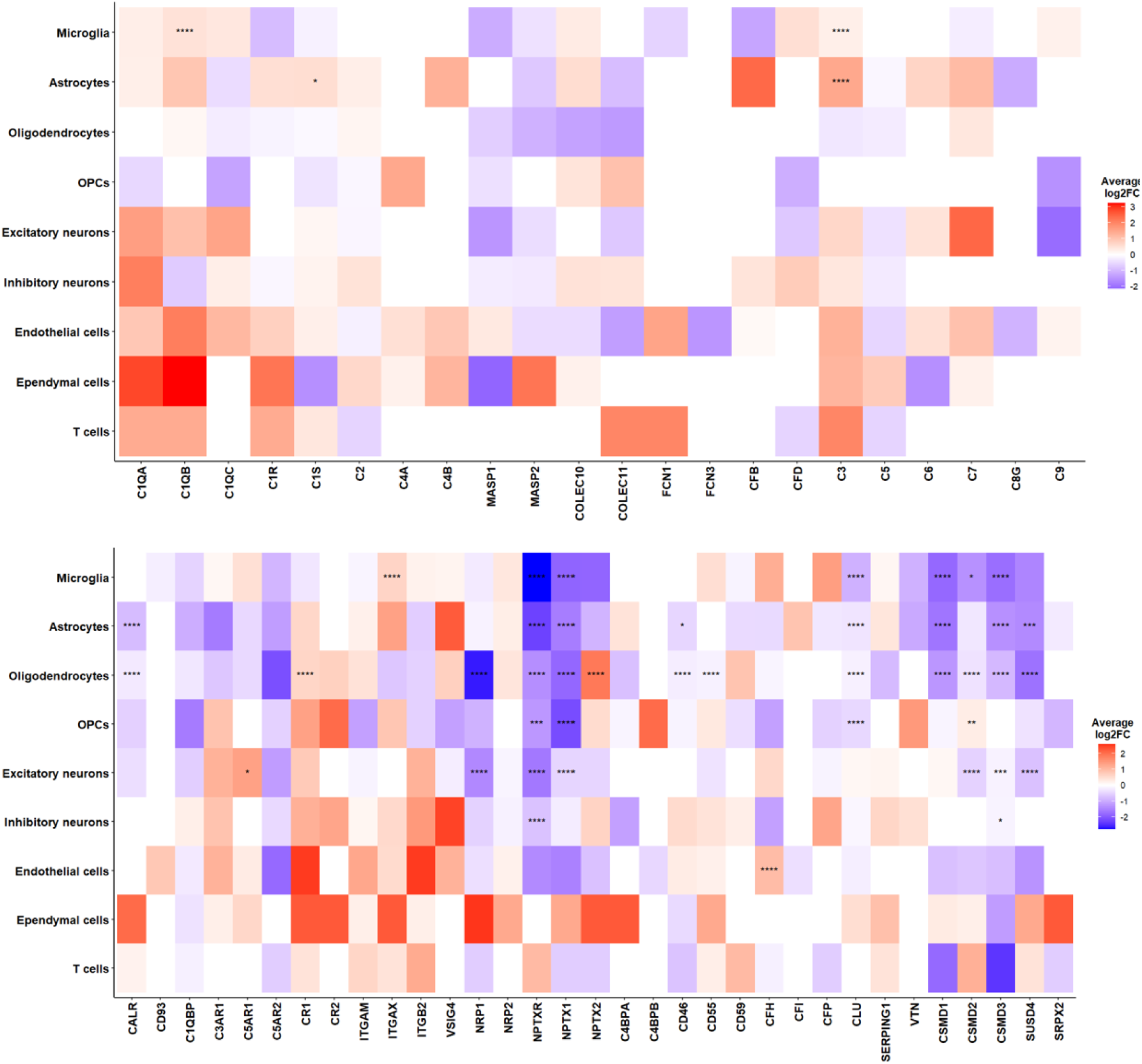
Differential expression of complement genes in AD cells compared to control cells. **A**. heatmap showing differences in complement gene expression in AD across cell types. Differential expression analysis (DEA) was run in every cell type, average log2 fold change is represented from low (blue) to high (red). A negative log2 fold change represents reduced gene expression in AD and a positive log2 fold change represents increased gene expression. Boxes show Bonferroni adjusted *P*-values (* = *P*<0.05, ** = *P*<0.01, *** = *P*<0.001, **** = *P*<0.0001). A. DEA summary for the genes of the complement pathways in the different cell types identified. *C1QB* was significantly upregulated in AD microglia and *C1S* was significantly upregulated in AD astrocytes. *C3* was significantly upregulated in AD microglia and astrocytes. **B**. DEA summary for complement receptor and regulator genes in the different cell types. Many cell types exhibited significant downregulation of genes encoding brain-associated putative complement genes in AD, including *NPTXR*, *NPTX1*, *CSMD1-3* and *SUSD4*. Some complement receptors were significantly upregulated in AD, notably *C5AR1* in excitatory neurons, *CR1* in oligodendrocytes and *ITGAX* in microglia.

**Supplementary Table 1.** Table of donor metadata used in the healthy hippocampus atlas.

| **Sample Number** | **orig.ident** | **dataset** | **Age** | **Sex** | **Donor ID** |
| --- | --- | --- | --- | --- | --- |
| 1 | SRR12905322 | GSE160189 | 26 | F | D1 |
| 2 | SRR12905324 | GSE160189 | 44 | M | D2 |
| 3 | SRR12905325 | GSE160189 | 24 | M | D3 |
| 4 | SRR12905326 | GSE160189 | 37 | F | D4 |
| 5 | SRR12905330 | GSE160189 | 24 | M | D3 |
| 6 | SRR12905331 | GSE160189 | 37 | F | D4 |
| 7 | GSM5005608 | GSE163737 | 67 | M | D5 |
| 8 | GSM5005609 | GSE163737 | 92 | M | D6 |
| 9 | GSM6280585 | GSE163737 | 85 | F | D7 |
| 10 | GSM6280586 | GSE163737 | 87 | M | D8 |
| 11 | GSM5348375 | GSE175814 | 69 | M | D9 |
| 12 | GSM5348377 | GSE175814 | 63 | M | D10 |
| 13 | SRR16928876 | GSE186538 | 79 | F | D11 |
| 14 | SRR16928877 | GSE186538 | 79 | F | D11 |
| 15 | SRR16928878 | GSE186538 | 79 | F | D11 |
| 16 | SRR16928879 | GSE186538 | 79 | F | D11 |
| 17 | SRR16928880 | GSE186538 | 79 | F | D11 |
| 18 | SRR16928881 | GSE186538 | 51 | M | D12 |
| 19 | SRR16928882 | GSE186538 | 51 | M | D12 |
| 20 | SRR16928883 | GSE186538 | 51 | M | D12 |
| 21 | SRR16928884 | GSE186538 | 51 | M | D12 |
| 22 | SRR16928885 | GSE186538 | 51 | M | D12 |
| 23 | SRR16928886 | GSE186538 | 50 | F | D13 |
| 24 | SRR16928887 | GSE186538 | 50 | F | D13 |
| 25 | SRR16928888 | GSE186538 | 50 | F | D13 |
| 26 | SRR16928889 | GSE186538 | 50 | F | D13 |
| 27 | SRR16928890 | GSE186538 | 50 | F | D13 |
| 28 | SRR16928891 | GSE186538 | 48 | M | D14 |
| 29 | SRR16928892 | GSE186538 | 48 | M | D14 |
| 30 | SRR16928893 | GSE186538 | 48 | M | D14 |
| 31 | SRR16928894 | GSE186538 | 48 | M | D14 |
| 32 | SRR16928895 | GSE186538 | 48 | M | D14 |
| 33 | SRR16928896 | GSE186538 | 48 | M | D14 |
| 34 | SRR16928897 | GSE186538 | 48 | M | D14 |
| 35 | SRR16928898 | GSE186538 | 44 | M | D15 |
| 36 | SRR16928900 | GSE186538 | 44 | M | D15 |
| 37 | SRR16928901 | GSE186538 | 44 | M | D15 |
| 38 | SRR16928902 | GSE186538 | 44 | M | D15 |
| 39 | SRR16928903 | GSE186538 | 44 | M | D15 |
| 40 | SRR16928904 | GSE186538 | 44 | M | D15 |
| 41 | SRR16928905 | GSE186538 | 48 | M | D16 |
| 42 | SRR16928906 | GSE186538 | 48 | M | D16 |
| 43 | SRR16928907 | GSE186538 | 48 | M | D16 |
| 44 | SRR16928908 | GSE186538 | 48 | M | D16 |
| 45 | SRR16928909 | GSE186538 | 48 | M | D16 |
| **46** | SRR16928910 | GSE186538 | 48 | M | D16 |
| **47** | SRR16928911 | GSE186538 | 48 | M | D16 |
| **48** | SRR16928912 | GSE186538 | 48 | M | D16 |

**Supplementary Table 2.** Table of donor metadata used in the mixed, healthy and AD hippocampus atlas.

| **Sample Number** | **orig.ident** | **Dataset** | **Condition** | **Age** | **Sex** | **Donor ID** | **Notes** |
| --- | --- | --- | --- | --- | --- | --- | --- |
| **1** | GSM4120422 | GSE138852 | AD | 89.4 | M | D1 | Two donors were combined into one sample in the original dataset |
| **1** | GSM4120422 | GSE138852 | AD | 83.8 | M | D2 | Two donors were combined into one sample in the original dataset |
| **2** | GSM4120423 | GSE138852 | AD | 73 | M | D3 | Two donors were combined into one sample in the original dataset |
| **2** | GSM4120423 | GSE138852 | AD | 74.6 | M | D4 | Two donors were combined into one sample in the original dataset |
| **3** | GSM4120424 | GSE138852 | AD | 67.8 | F | D5 | Two donors were combined into one sample in the original dataset |
| **3** | GSM4120424 | GSE138852 | AD | 83 | F | D6 | Two donors were combined into one sample in the original dataset |
| **4** | GSM4120425 | GSE138852 | Control | 77.5 | M | D7 | Two donors were combined into one sample in the original dataset |
| **4** | GSM4120425 | GSE138852 | Control | 82.7 | M | D8 | Two donors were combined into one sample in the original dataset |
| **5** | GSM4120426 | GSE138852 | Control | 72.6 | M | D9 | Two donors were combined into one sample in the original dataset |
| **5** | GSM4120426 | GSE138852 | Control | 75.6 | M | D10 | Two donors were combined into one sample in the original dataset |
| **6** | GSM4120428 | GSE138852 | Control | 90.8 | M | D11 | Two donors were combined into one sample in the original dataset |
| 6 | GSM4120428 | GSE138852 | Control | 84.7 | M | D12 | Two donors were combined into one sample in the original dataset |
| 7 | GSM4120429 | GSE138852 | AD | 91 | M | D13 | Two donors were combined into one sample in the original dataset |
| 7 | GSM4120429 | GSE138852 | AD | 83.8 | M | D14 | Two donors were combined into one sample in the original dataset |
| 8 | GSM5348374_A1 | GSE175814 | AD | Unclear | Unclear | D15 | Original authors cite donors being Braak stage III/IV at GSE175814 but Braak stage I/II in their paper; PMID: 36827281. |
| 9 | GSM5348375_A2 | GSE175814 | Control | 69 | M | D16 |  |
| 10 | GSM5348376_A3 | GSE175814 | AD | Unclear | Unclear | D17 | Original authors cite donors being Braak stage III/IV at GSE175814 but Braak stage I/II in their paper; PMID: 36827281. |
| 11 | GSM5348377_A4 | GSE175814 | Control | 63 | M | D18 |  |
| 12 | WT_1304 | GSE212606 | Control | 81 | M | D19 |  |
| 13 | WT_5459 | GSE212606 | Control | 70 | F | D20 |  |
| 14 | WT_1306 | GSE212606 | Control | 83 | F | D21 |  |
| 15 | AD_1248 | GSE212606 | AD | 86 | F | D22 |  |
| 16 | AD_5325 | GSE212606 | AD | 93 | F | D23 |  |
| 17 | WT_5356 | GSE212606 | Control | 94 | M | D24 |  |
| 18 | WT_1247 | GSE212606 | Control | 94 | M | D25 |  |
| 19 | AD_1282 | GSE212606 | AD | 89 | M | D26 |  |
| 20 | AD_1291 | GSE212606 | AD | 83 | M | D27 |  |
| 21 | WT_1311 | GSE212606 | Control | 85 | F | D28 |  |
| 22 | AD_1316 | GSE212606 | AD | 89 | F | D29 |  |
| 23 | AD_5245 | GSE212606 | AD | 80 | M | D30 |  |

*This is an original research manuscript currently under consideration for publication in Brain. It has not been published elsewhere and is being made available as a preprint to promote open access and early dissemination of our findings*.

